# Moderating Effect of Social Support on Role Conflict and Job Satisfaction among Nurses with Multiple Roles

**DOI:** 10.1101/2020.01.14.906263

**Authors:** Caifu Li, Rhayun Song, Xing Fan, Lijuan Xu

## Abstract

Registered nurses perform multiple roles in their lives at work, at home, and at school when they decide to pursue various types of advanced degrees. Work-family conflict was found to be negatively correlated with job satisfaction among nurses. This study aimed to determine the moderating effect of social support on the relationship between role conflicts and job satisfaction among registered nurses in mainland China. The study had a cross-sectional correlational design. Convenience sampling was applied to recruit 320 nurses with multiple roles from 8 nursing universities and 3 hospitals in China between January and August 2017. SPSS (version 20.0) software and PROCESS program was used to examine to moderating effect of social support between the relationship of role conflicts and job satisfaction. It was found that role conflicts were negatively correlated with job satisfaction. Support from work and support from family negatively moderated the relationship between role conflicts and job satisfaction. These findings suggest that support from family, coworkers and nursing administrators, and the implementation of school- and family-friendly policies would help nurses who are pursuing further nursing degrees to handle their multiple roles and improve their job satisfaction.

## INTRODUCTION

Learning new knowledge and pursuing education are very important to improving career prospects. There are three types of educational programs in China for training registered nurses (RNs): secondary nursing education programs, associate-degree nursing programs, and bachelor-degree nursing programs. Approximately one-quarter (22.8%) of Chinese RNs had a secondary nursing degree in 2015 [1]. The Chinese National Plan for the Development of Nursing Career implemented a policy of improving the education of RNs in 2005, which has led to increases in the number of RNs with secondary nursing degrees participating in RN-BSN education programs [2]. Moreover, nursing science has been authorized a first-level discipline in 2011, which has further increased the number of RNs pursing postgraduate degrees in mainland China [3].

RNs pursuing further nursing degrees assume a student role that is additional to the roles in their family and their professional career. An RN marrying and deciding to pursue a further nursing degree is likely to experience role conflicts stemming from the need to perform these multiple roles. Kahn et al. defined role conflicts as stemming from the difficulty of addressing multiple pressures that occur simultaneously [4]. Work-family-school role conflicts were forms of interrole conflicts in which pressures from the work, family, and school domains are mutually incompatible, including work and family roles interfere with school roles, school and family roles interfere with work roles, and work and school roles interfere with family roles [5]. Performing a larger number of roles will increase the likelihood of the individual experiencing role conflicts and negative outcomes, including decreased job dissatisfaction.

### Role conflict and job satisfaction

Job satisfaction is influenced by the emotions and feelings that a person has about working [6]. Since the job satisfaction of nurses is correlated with their retention and performance, there is a need to quantify job satisfaction and its influencing factors among RNs. The association between work-family conflict and job satisfaction was well cited in nursing literature [7-9]. Reducing conflict between family and work roles can improve the quality and productivity of nursing, which will in turn improve job satisfaction [8, 10]. Both work-to-family and family-to-work role conflicts were found to be negatively correlated with job satisfaction, with work-to-family role conflict having a greater affect than family-to-work role conflict on job satisfaction among RNs in mainland China [7, 9]. There was few studies to describe the relationship between role conflicts and job satisfaction among RNs pursuing for further nursing degree.

### Moderating role of social support

Social support was referred as the emotional, informational, instrumental, and appraisal support a person received from others [11]. Role conflict specific social support was the degree to which individuals accept that others “care about their ability to experience role conflict and demonstrate this care by providing helpful social interaction and resources” [12]. Therefore, work-family-school specific social support could come from either the work domain, the family domain or the school domain. In the Social Support Buffering Hypothesis, social support serves as a protector that can induce an emotion (e.g., job satisfaction) before an individual experiences stressful event [13]. In this situation, social support could be treated as a moderator, individuals who receive more social support from others making the role conflict more manageable and less threatening might therefore be less likely to experience negative emotions. This Hypothesis was supported by the recently Job Demand Resources Model, which indicated that individuals’ resources could buffer the impact of initial stress on consequent health outcome [14].

Previous studies have shown that social support act as a moderator in the association of work-family conflict and psychological health [15-16]. In a study among Chinese doctors, Hao and colleagues demonstrated that perceived organizational support successfully moderated on the relationship between work-to-family conflict and depression, while organizational support had no moderating effect between family-to-work conflict and depression [15]. In another study among RNs, it was reported that support from friends and support from coworkers weakened the association between family-to-work conflict and job performance [16]. In a study about work-family-school role conflicts, social support was treated as a mediator between the relationship of work-family-school role conflicts and mental health [17]. It was unclear whether social support moderate the relationship of role conflicts and job satisfaction among RNs pursuing further nursing degree, and according to role conflict specific social support perspective, which kind of social support was most effective to lesson the effect of role conflict on job satisfaction among RNs pursuing further nursing degree was not clear.

Other studies have identified that job satisfaction is associated with age [18-19], marital status [20-21], clinical experience [19], educational level [18, 22], and work patterns [19, 23].

### Hypothesis

Based on a review of the literature and assumptions inherent in the Social Support Buffering Hypothesis and Job Demand Resources Model, we stated the following hypotheses:

Hypothesis 1: Role conflicts would negatively correlate with job satisfaction.

Hypothesis 2: Social support would moderates the relationship between role conflicts and job satisfaction; more specifically, (a) support from family moderates the relationship between role conflicts and job satisfaction; (b) support from work moderates the relationship between role conflicts and job satisfaction; and (c) support from school moderates the relationship between role conflicts and job satisfaction.

## METHODS

### Design

This study employed a cross-sectional correlational design to examine the relationship between role conflicts and job satisfaction, and explore the moderating effect of social support on the relationship between role conflicts and job satisfaction.

### Participants

The participants were a convenience sample of 320 RNs currently employed and/or enrolled at 8 nursing universities and 3 hospitals in China. The inclusion criteria for the subjects were (1) currently enrolled in advanced degree programs, (2) currently working as a nurse for one or more years.

### Data collection

The study obtained data using a voluntary paper-based survey. A single contact person in each hospital or university was asked to distribute and collect the questionnaires from January to August 2017.

### Instruments

#### Role conflicts

The work–family–school role conflict scale developed by Xu and Song [5] was used to measure role conflicts in the present study. This scale comprises 10 items in the following 3 dimensions: family–work-to-school, family–school-to-work, and work–school-to-family role conflicts. The scale reportedly has good validity, with a reliability coefficient of 0.82 [24]. The original scale was translated in the present study into Chinese by three Korean-Chinese nursing researchers using a forward-and-back-translation method.

The internal consistencies for family–work-to-school, family–school-to-work, and work–school-to-family role conflicts were confirmed in the present study by the internal-consistency coefficients having values of 0.83, 0.75, and 0.70, respectively. The construct validity of work–family–school role conflicts was examined using confirmatory factor analysis (CFA), which yielded the following results for the scale: comparative fit index (CFI)=0.96, goodness-of-fit index (GFI)=0.95, root-mean-square error of approximation (RMSEA)=0.061, 90% confidence interval (CI) of RMSEA=0.055–0.079, standardized root-mean-square residual (SRMR)=0.054, and c^2^/df (degrees of freedom)=2.15 (p<0.001).

#### Role-related social support

The role-related social support scale developed by Xu and Song [5] was used to measure social support in this study. This scale comprises nine items in three dimensions (social support from friends, social support from work, and social support from family), and each item is scored from 1 (“strongly disagree”) to 5 (“strongly agree”). Goong et al. [24] reported that this scale has good construct validity, with an internal reliability coefficient of 0.75. The scale was translated in the present study into Chinese by three Korean-Chinese nursing researchers using a forward-and-back-translation method.

The presence of internal consistency for social support from school, social support from family, and for social support from work in the present study was confirmed by the internal-consistency coefficients having values of 0.70, 0.63, and 0.63, respectively. CFA yielded the following parameters for the scale: CFI=0.98, GFI=0.97, RMSEA=0.012, 90% CI of RMSEA=0.0–0.049, SRMR=0.035, and c^2^/df=1.31 (p=0.089).

#### Job satisfaction

The Andrews and Withey job satisfaction questionnaire [25] was used to measure general job satisfaction. This scale showed a reliability coefficient of 0.83 in a previous study [26]. In the present study the internal-consistency coefficient of this questionnaire was 0.86. CFA produced the following parameters for the questionnaire: CFI=0.99, GFI=0.99, RMSEA=0.031, 90% CI of RMSEA=0.019–0.059, SRMR=0.013, and c^2^/df=1.29 (p=0.272).

### Ethical considerations

This study was approved by the Institutional Review Board at Medical and Health College of Lishui University (No. 20160124). Potential study participants were provided with an explanation of the study purpose and methods, as well as assurances about their privacy, confidentiality, and right to withdraw from the study without penalty.

### Data analysis

Data were analyzed using the SPSS (version 20.0) software and PROCESS program. The demographic characteristics and measured variables were analyzed using descriptive statistics. Relationships among the measured variables were quantified using Pearson correlation coefficients. The effect of role conflicts on job satisfaction, and the moderating effect of social support between the relationship of role conflicts and job satisfaction was examined using PROCESS program.

## RESULTS

### Descriptive results

The demographic characteristics of the participants were described in Table 1. They were aged 27.47±6.88 years (mean±SD), and had an age range from 20 to 49 years. Approximately 40% of the subjects were married, 59.4% were single, and 30% of them had from one to two children. Their length of clinical experience was 5.95±6.45 years. Most (75.6%) of the subjects were staff nurses, with the remaining 24.4% being nursing managers or charge nurses. Most of the participants worked day, evening, and night shifts (61.9%), with the other 38.1% working day shifts only. There were only 15.3% participants enrolled in a doctoral or master’s program and 69.7% enrolled in an RN-BSN program.

**Table 1.** Demographic characteristics of the study subjects.

| Characteristic | Categories | Value<sup>†</sup> |
| --- | --- | --- |
| Age, years |  | 27.47±6.88 |
| Clinical experience, years |  | 5.95±6.45 |
| Marital status | Married | 125 (39.1%) |
|  | Single | 190 (59.4%) |
|  | others | 5(1.6%) |
| Have children | Yes | 97 (30.3%) |
|  | No | 223 (69.7%) |
| Work position | Staff nurse | 242 (75.6%) |
|  | Charge nurse or higher | 78 (24.4%) |
| Work pattern | Fixed day shifts | 122 (38.1%) |
|  | With night shift | 198 (61.9%) |
| Enrolled program | Master's or doctoral program | 49 (15.3%) |
|  | RN-BSN program | 271 (84.7%) |

Note: <sup>†</sup> Values are mean±SD or n (%) .

The mean and SD for role conflicts, social support and job satisfaction are listed in Table 2. The level of overall role conflicts was assessed as being moderate to high, with a score of 3.21±0.49. The respondents generally perceived that they received favorable levels of social support, with a score of 3.39±0.42. Job satisfaction was assessed as favorable, with a score of 3.08±0.59.

**Table 2.** Levels of role conflicts, social support, and job satisfaction, and relationships among all the variables.

| Variable | Mean ± SD | 1 | 2 | 3 | 4 | 5 | 6 | 7 | 8 | 9 | 10 |
| --- | --- | --- | --- | --- | --- | --- | --- | --- | --- | --- | --- |
| Age, years (1) |  | -- |  |  |  |  |  |  |  |  |  |
| Clinical experience, years (2) |  | 0.88** | -- |  |  |  |  |  |  |  |  |
| Getting married (3) |  | 0.60** | 0.58** | -- |  |  |  |  |  |  |  |
| Having children (4) |  | 0.68** | 0.67** | 0.80** | -- |  |  |  |  |  |  |
| Staff nurses (5) |  | 0.59** | 0.61** | 0.40** | 0.45** | -- |  |  |  |  |  |
| With night shift (6) |  | -0.19* | -0.21** | -0.18* | -0.22** | -0.13* | -- |  |  |  |  |
| RN-BSN program (7) |  | 0.32** | -0.20** | -0.25** | 0.24* | 0.24** | -0.10 | -- |  |  |  |
| Role conflicts (8) | 3.21±0.49 | 0.15* | 0.16* | 0.17* | 0.12* | 0.18* | -0.07 | 0.08 | -- |  |  |
| Social support (9) | 3.39±0.42 | -0.03 | 0.02 | 0.13* | 0.06 | 0.12* | 0.10 | 0.04 | -0.26** | -- |  |
| Job satisfaction (10) | 3.08±0.59 | .014* | 0.15* | 0.19* | 0.18* | 0.11 | -0.16* | -0.02 | -0.24** | 0.28** | -- |

Note: \*p<0.05; \*\*p<0.001

### Test for collinearity

From table 2, except for work position (staff nurses) and Enrolled education program (RN-BSN program), all other demographic variables were significant correlated with job satisfaction (p<0.05). Among those variables that have significant relationship with job satisfaction, age had a high relationship with clinical experience (r=0.88, p<0.001), and getting married had a high relationship with having children (r=0.80, p<0.001). Therefore, only clinical experience, having children, and with night shift was included as covariate variables in the following analysis of testing the moderating effect of social support between relationship of role conflicts and job satisfaction.

### Moderating effect

As shown in table 3, after controlling the covariate variables (work experience, having children, with night shift), role conflicts was negatively correlated with job satisfaction (β=-0.19, p=0.006). The interaction of conflicts and social support were significantly associated with job satisfaction (β=0.18, p=0.040). In addition, conditional effect in the PROCESS analysis, showed that when the moderator (social support) is below the average (average =0), there was a significant negative relationship between conflicts and job satisfaction (β=-0.24 p<0.001); when the moderator (support from family) is equal (β=-0.17, p<0.001) and higher (β=-0.11, p=0.015) than the average, a negative relationship between role conflicts and job satisfaction became smaller yet significant (Figure1).

**Figure 1.**
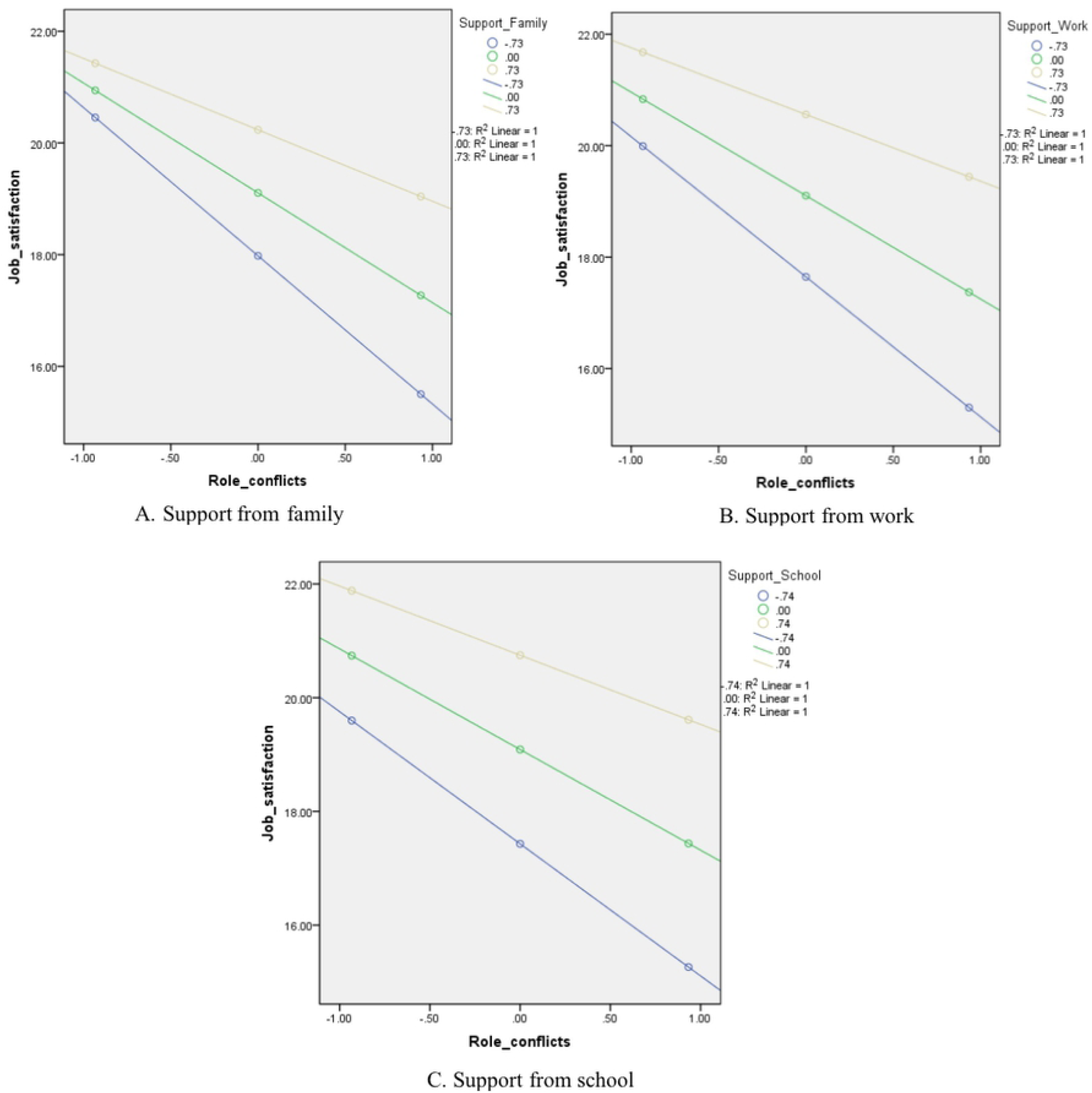
Moderating role of social support on the relationship between role conflicts and job satisfaction

**Table 3.** Role conflicts as the independent variable and social support as the moderator.

| Variables | $\beta$ | t | p | R<sup>2</sup> |
| --- | --- | --- | --- | --- |
| Work experiences | 0.03 | 0.05 | 0.954 | 0.16 |
| Having children | 0.11 | 1.67 | 0.100 |  |
| With night shift | -0.09 | -1.03 | 0.305 |  |
| Role conflicts | -0.19 | -3.13 | 0.006 |  |
| Social support | 0.28 | 4.11 | <0.001 |  |
| Interaction | 0.18 | 2.06 | 0.040 |  |

Note: $\Delta R^2$ (due to interaction)= 0.017 (p=0.040).

As shown in table 4 the interaction of conflicts and support from family were significantly associated with job satisfaction (β=0.21, p=0.028). In addition, conditional effect in the PROCESS analysis, showed that when the moderator (support from family) is below the average (average =0), there was a significant negative relationship between conflicts and job satisfaction (β=-0.30 p<0.001); when the moderator (support from family) is equal (β=-0.19, p<0.001) and higher (β=-0.12, p=0.039) than the average, a negative relationship between role conflicts and job satisfaction became smaller yet significant (Figure1).

**Table 4.** Moderating effect of social support with sub-dimensions of support.

| Variables | $\beta$ | t | p | R<sup>2</sup> |
| --- | --- | --- | --- | --- |
| Support from family |  |  |  |  |
| Work experiences | 0.03 | 0.05 | 0.954 | 0.17 |
| Having children | 0.11 | 1.67 | 0.100 |  |
| With night shift | -0.09 | -1.03 | 0.305 |  |
| Role conflicts | -0.20 | -2.01 | 0.031 |  |
| Support from family | 0.34 | 4.24 | <0.001 |  |
| Interaction | 0.21 | 2.02 | 0.028 |  |
| $\Delta R^2(\text{due to interaction})= 0.022$ (p=0.028). | | | | |
| Support from work |  |  |  |  |
| Work experiences | 0.03 | 0.25 | 0.806 | 0.15 |
| Having children | 0.11 | 1.47 | 0.146 |  |
| With night shift | -0.08 | -0.48 | 0.632 |  |
| Role conflicts | -0.22 | -2.02 | 0.002 |  |
| Support from work | 0.31 | 3.57 | <0.001 |  |
| Interaction | 0.22 | 2.04 | 0.038 |  |
| $\Delta R^2(\text{due to interaction})= 0.020$ (p=0.038). | | | | |
| Support from School |  |  |  |  |
| Work experiences | 0.02 | 0.06 | 0.959 | 0.16 |
| Having children | 0.10 | 1.42 | 0.100 |  |
| With night shift | -0.08 | -0.89 | 0.545 |  |
| Role conflicts | -0.22 | -2.29 | <0.001 |  |
| Support from school | 0.30 | 3.68 | <0.001 |  |
| Interaction | 0.18 | 1.13 | 0.090 |  |
| $\Delta R^2(\text{due to interaction})= 0.004$ (p=0.258). | | | | |

As shown in table 4 the interaction of role conflicts and support from work were significantly associated with job satisfaction (β=0.22, p=0.038). In addition, conditional effect in the PROCESS analysis, showed that when the moderator (support from work) is below the average (average =0), there was a significant negative relationship between role conflicts and job satisfaction (β=-0.29, p<0.001); when the moderator (support from work) is equal the average, a negative relationship between role conflicts and job satisfaction became smaller yet significant (β=-0.20, p<0.001) (Figure 1).

As shown in table 4, the interaction of role conflicts and support from school were not associated with job satisfaction (β=0.18, p=0.090) (Figure1).

## DISCUSSION

This study aimed to elucidate the effects of role conflicts on job satisfaction, as moderated by social support. The obtained results support the proposed hypotheses, with social support from family and work found to negatively moderated the relationship between role conflicts and job satisfaction among RNs pursuing further nursing degree.

There are several theoretical and practical implications of these findings. Firstly, role conflicts was a serious problems among RNs pursuing further nursing degree in mainland China, and is consistent with the previous studies [24] which showing that RNs enrolled in educational programs got into trouble to deal with role conflicts. Females have traditionally been responsible for family affairs in Asian countries, and so when they also adopt a worker or student role this would lead to the potential for conflict among these multiple domains. That is, in addition to their education, female nurses are likely to maintain the responsibilities they have in their work and family, which means that role conflicts are a significant challenge to RNs who enroll in an education program.

Secondly, role conflicts was found to be negatively correlated with job satisfaction, which is consistent with previous reports [9, 10, 21]. The present study further demonstrated the presence of work–family–school role conflicts in RNs enrolled in a nursing education program. Among the dimensions of role conflicts, the score was highest for work–school-to-family role conflict, which also had the greatest influence on job satisfaction. There have been similar reports of job satisfaction being influenced more by work-to-family role conflict than by family-to-work role conflict, which is due to the high intensity of nursing work and the associated high risks of adverse outcomes and irregular working hours [9, 27].

Thirdly, support from family and work were found to moderate the relationship between role conflicts and job satisfaction, and especially support from work was the most effective to lessen role conflicts, and increase their job satisfaction. For the nurses with a low level of support from work, job satisfaction will be decreased with the increase of role conflicts. This supports Job Demand Resources Model [14], which indicated that individuals’ resources could buffer the impact of initial stress on consequent health outcome, and is consistent with Nohe & Sonntag [28] showing that leader support buffered the relationship between work-family conflict and turnover intentions. Cortese et al. [27] suggested that the job satisfaction of nurses could decrease when there is a less support from management, with this relationship being weaker in the presence of work–family role conflict. In another study, it was also reported that coworker support moderated the relationship between work-family conflict and job performance among RNs [16]. In the present study, it was shown that coworker support, supervisor support and organizational support was the most effective kind of social support to make nurses feel more satisfied, and staying in their job.

### Practical implications

The findings of this study have practical implications for both nursing managers and organizations. Managers should attempt to facilitate the social support that RNs enrolled in nursing education programs receive from the hospital organization and family, since this can both decrease their role conflicts and increase their job satisfaction. Firstly, hospital organizations could implement family-welfare policies for RNs, such as running nurseries, providing paid personal leave, and making policy of caring for the elderly of the staffs. Secondly, nursing managers should encourage RNs to go on to further nursing education, and implement school-friendly policies such as paying tuition fees and implementing flexible work schedules. Thirdly, nursing administrators should offer emotional support and provide a work atmosphere that is conducive to helping RNs to handle their role conflicts. Such strategies could also be valuable in encouraging more RNs to pursue further nursing degrees.

### Limitations

One of the limitations of this study was its cross-sectional design, which prevents assessments of changes and trends over time and so yields weaker evidence for causal relationships among the study variables. The second limitation involved the utilization of convenience sampling, which might have resulted in the included sample not being representative of the target population. Although this would compromise the generalizability of the research findings, the possibility of problem was minimized by the sample in the present study being quite diverse in terms of marital status, education, work patterns, clinical experience, and age.

## CONCLUSION

This study has revealed that there are significant relationships between role conflicts and job satisfaction among RNs pursuing further nursing degree. Support from family and work was found to negatively moderate the relationship between role conflicts and job satisfaction among RNs pursuing further nursing degree. The findings of this study provide valuable information that could be used when formulating new policies related to nursing. RNs enrolled in nursing education programs exhibit role conflicts that contribute significantly to job dissatisfaction. These RNs need to receive social support from organizations, supervisors, and family in order to relieve the role conflicts and increase their job satisfaction.

## Supplementary Materials

## Author Contributions

Conceptualization, Caifu Li and Hwasoo Koong; Methodology, Lijuan Xu and Rhayun Song; Formal Analysis, Rhayun Song; Investigation, Xing Fan; Data Curation, Lijuan Xu; Writing –Original Draft Preparation, Lijuan Xu; Writing – Review & Editing, Hwasoo Koong; Supervision, Hwasoo Koong; Project Administration, Caifu Li; Funding Acquisition, Caifu Li”

## Funding

This research was supported by the Research Foundation of Lishui city--2017RKX09.

## Acknowledgments

The authors thank all the universities, hospitals, and nurses for their participants.

## Conflicts of Interest

The authors declare no conflict of interests.

## REFERENCES

[1] NHFPCPRC (National Health and Family Planning Commission of the People’s Republic of China). “The National Plan for the Development of Nursing Career (2016–2020)”, March 20, 2017, URL:http://www.moh.gov.cn/yzygj/s3593/201611/92b2e8f8cc644a899e9d0fd572aefef3.shtml.

[2] MHPRC (Ministry of Health of the People’s Republic of China). “Outline of the National Plan for the Development of Nursing Career (2005–2010)”, March 20, 2017, URL: http://www.110.com/fagui/law_112370.html.

[3] MEPRC (Ministry of Education of the People’s Republic of China). “Notification about the List of First-level Discipline with Scientific System added in 2010”, March 15, 2017, URL: http://www.moe.gov.cn/srcsite/A22/yjss_xwgl/moe_818/201103/t20110303_117375.html.

[4] Kahn, R. L.; Wolfe, D. M.; Quinn, R. P.; Snoek, J. D. Organizational Stress: Studies in Role Conflict and Ambiguity; John Wiley & Sons: New York, USA, 1964; PP 378–397.

[5] Xu, L.; Song, R. Development and validation of the work-family-school role conflicts and role-related social support scales among registered nurses with multiple roles. International Journal of Nursing Studies. 2013, 50(10), 1391–1398. DOI: 10.1016/j.ijnurstu.2013.01.003.

[6] Spector, P. E. Job satisfaction: Application, Assessment, Causes, and Consequences; SAGE: London, England, 1997; PP 124-156.

[7] AlAzzam, M; AbuAlRub, R.F.; Nazzal, A.H. The Relationship Between Work-Family Conflict and Job Satisfaction Among Hospital Nurses. Nursing Forum. 2017, 52(4), 278–288. doi: 10.1111/nuf.12199.

[8] Ding, X.; Yang, Y.; Su, D.; Zhang, T.; Li, L,.; Li, H. Can Job Control Ameliorate Work-family Conflict and Enhance Job Satisfaction among Chinese Registered Nurses? A Mediation Model. The International Journal of Occupational and Environmental Medicine. 2018, 9(2), 97–105.

[9] Zhao, S.; Chen, H. Research progress on nurses’ work family conflict and job satisfaction. Chinese Nursing Research. 2015, 29(2C), 644–646. DOI: 10.3969/j.issn.1009-6493.2015.06.002.

[10] Zhou, H.; Chang, H.; Liu, D.; Guo, J. Correlation of work-family conflict, social support and job satisfaction of nurses. Chinese Nursing Management. 2011, 11(10), 57–60. doi: 10.15171/ijoem.2018.1176.

[11] House, J. S. Job stress and social support; Reading, MA: Addison-Wesley: USA, 1981.

[12] Kossek, E. E.; Pichler, S.; Bodner, T.; & Hammer, L. B. Workplace social support and work– family conflict: A meta-analysis clarifying the influence of general and work–family-specific supervisor and organization support. Personnel Psychology. 2011, 64(2), 289–313. http://dx.doi.org/10.1111/j.1744-6570.2011.01211.x.

[13] Bakker, A.B.; Demerouti, E.; Euwema, M.C. Job resources buffer the impact of job demands on burnout. Journal of Occupational Health Psychology. 2005, 10(2), 170–80.

[14] Cohen, S.; Wills, T. A. Stress, social support, and the buffering Hypothesis. Psychological Bulletin. 1985, 98(2):310–357.

[15] Hao, J.; Wang, J.; Liu, L.; Wu, W.; Wu, H. Perceived Organizational Support Impacts on the Associations of Work-Family Conflict or Family-Work Conflict with Depressive Symptoms among Chinese Doctors. International Journal of Environmental Research and Public Health. 2016, 13(3), pii: E326. doi: 10.3390/ijerph13030326.

[16] Wang, M.L.; Tsai. L.J. Work-family conflict and job performance in nurses: the moderating effects of social support. The Journal of Nursing Research. 2014, 22(3), 200–207. doi: 10.1097/jnr.0000000000000040.

[17] Xu, L.; Song, R. Influence of work-family-school role conflicts and social support on psychological wellbeing among registered nurses pursuing advanced degree. Applied Nursing Research,. 2016, 31, 6–12. DOI: 10.1016/j.apnr.2015.12.005.

[18] Al Maqbali, M. A. Factors that influence nurses’ job satisfaction: a literature review. Nursing Management. 2015, 22(2), 30–37. DOI: 10.7748/nm.22.2.30.e1297.

[19] Wan, X.; Wang, Y.; Yu, H.; Liu, L. Status quo of work-family and family-work conflicts in nurses in operating room. Modern Clinical Nursing. 2017, 16(3), 52–56. DOI: 10.3969/j.issn.1671-8283.2017.03.014.

[20] Ghawadra, S.F.; Abdullah, K.L.; Choo, W.Y.; Phang, C.K. Psychological distress and its asociation with job satisfaction among nurses in a teaching hospital. Journal of Clinical Nursing. 2019, doi: 10.1111/jocn.14993.

[21] Gao, Y.; Shi, J.; Niu, Q.; Wang, L. Work-family conflict and job satisfaction: emotional sintelligence as a moderator. Stress and Health. 2012, 29(3), 222–228. DOI: 10.1002/smi.2451.

[22] Nguyen, H.V.; Duong, H.T; Vu, T.T. Factors associated with job satisfaction among district hospital health workers in Northern Vietnam: a cross-sectional study. International Journal of Health Planning and Management. 2017, 32(2), 163–179. doi: 10.1002/hpm.2337.

[23] Aloisio, L.D.; Gifford, W.A.; McGilton, K.S.; Lalonde, M.; Estabrooks, C.A.; Squires, J.E. Factors Associated With Nurses’ Job Satisfaction In Residential Long-term Care: The Importance of Organizational Context. Journal of American Medical Directors Association. 2019, *pii: S1525-8610*(19), 30515–8. doi: 10.1016/j.jamda.2019.06.020.

[24] Goong, H.; Xu, L.; & Li, C. Effect of work–family–school role conflicts and role-related social support on burnout in registered nurses: a structural equation modeling approach. Journal of Advanced Nursing. 2016, 72(11), 2762–2772. DOI: 10.1111/jan.13029.

[25] Rentsch, J. R.; Steel, R. P. Construct and concurrent validation of the Andrews and Withey job satisfaction questionnaire. Education and Psychological Measurement. 1992, 52(2), 357–367. DOI: 10.1177/0013164492052002011.

[26] van Sanne, N.; Sluiter, J. K.; Verbeek, J. H. A. M.; Frings-Dresen, M. H. W. Reliability and validity of instruments measuring job satisfaction—a systematic review. Occupational Medicine. 2003, 53(3), 191–200. DOI: 10.1093/occmed/kqg038.

[27] Cortese, C. G.; Colombo, L.; Ghislieri, C. Determinants of nurses’ job satisfaction: the role of work-family conflict, job demand, emotional charge and social support. Journal of Nursing Management. 2010, 18(1), 35–43. DOI: 10.1111/j.1365-2834.2009.01064.x.

[28] Nohe, C.; Sonntag, K. Work–family conflict, social support, and turnover intentions: A longitudinal study. Journal of Vocational Behavior. 2014, 85, 1–12. doi:org/10.1016/j.jvb.2014.03.007.

